# Pseudo-observations of longevity improve detection of resilience-associated genes in the Radiation Late Effects Cohort of rhesus macaques

**DOI:** 10.64898/2026.11.02.763827

**Authors:** Juliana H. Lawandos, Mitchell Paukner, George W. Schaaf, John D. Olson, Muthuswamy Raveendran, R. Alan Harris, Jeffrey Rogers, J. Mark Cline, Ellen E. Quillen

## Abstract

Non-human primates are an ideal model for interrogating the genetic basis of human lifespan and resilience due to their long lifespan compared to other model organisms, development of similar age-related chronic conditions, and genetic similarity. However, like humans, statistical methods for modeling survival suffer from the challenges of dealing with non-proportionality of hazards, large numbers of living individuals that must be censored, and loss-to-follow-up over studies that may last a decade or more. Here we overcome these challenges through the use of pseudo-observations of Restricted Mean Survival Time as a measure of resilience to the acute challenge of radiation exposure and chronic challenges of aging and a Western-style diet in rhesus macaques (*Macaca mulatta*) from the Wake Forest Non-Human Primate Radiation Late Effects Cohort. Our whole-exome analysis used Genome-wide Efficient Mixed Model Association (GEMMA) to identify variants associated with longevity, and a subsequent Gene Set Enrichment Analysis (GSEA) highlighted the cadherin and integrin signaling pathways as central to this resilience. As dysregulation of cell adhesion and extracellular matrix signaling is central to the development of fibrosis, we tested the functional relevance of these signals using a targeted Combined Annotation Dependent Depletion (CADD) score-weighted Sequence Kernel Association Test-Optimal (SKAT-O) analysis of GSEA leading-edge genes from both the cadherin and integrin pathways. This linked variant burden in *PCDHGA5, PCDHGA3, PCDHGB2, PCDHA11,* and *PSENEN* to the severity of myocardial fibrosis, and in *IBSP, PDGFD,* and *LAMC2* to the severity of pulmonary fibrosis observed at necropsy. Critically, we note that the genetic variants are not caused by the radiation but rather create an underlying risk for disease that is then exacerbated by the acute and persistent stressors challenging these animals’ intrinsic resilience. The success of this method in identifying biologically relevant genetic variants underlying survival in animals undergoing a strong challenge to their resilience highlights not only the role of genes predisposing animals to fibrosis in lifespan, but the overall utility of pseudo-observations in overcoming the limitations of survival analyses in humans and other long-lived species.

---

Longevity is the result of surviving a lifetime of environmental challenges through intrinsic resilience. It follows then that capturing the genetic factors contributing to this intrinsic resilience is readily accomplished in the context of extreme challenges to resilience. Here, we consider the Wake Forest Radiation Late Effects Cohort, a cohort of rhesus macaques (*Macaca mulatta)* that have undergone the unique combination of an acute insult to resilience from early-life irradiation and the chronic stressors of aging and a high-fat diet. Despite these environmental challenges, many animals in the cohort have lifespans similar to those seen in non-irradiated animals, and there is substantial variability in lifespan that is not explained by radiation dose, suggesting a biological basis for resilience (Quillen et al., 2025). We have previously described the utility of this cohort for understanding the underlying biology of aging through the lens of resilience/vulnerability to stressors – a characteristic of increased risk of mortality among older adults (Schaaf et al., 2023).

Recent analyses of the heritability of human lifespan, as well as animal models, suggest more than 50% of the variance in lifespan is attributable to genetics (Quillen et al., 2025). However, studies of genetic contributors to lifespan face several statistical and logistical challenges among humans and our nearest relatives, non-human primates. In all species, traditional survival analysis techniques may not be appropriate for studying lifespan, as risk of death is not evenly distributed across ages (i.e. the hazards are not proportional throughout observation). In animal models, lifespan is difficult to measure due to the ethical imperative of euthanizing animals to minimize pain associated with end-of-life disease, as well as interventional studies which may necessitate sacrifice of the animal prior to the end of their healthspan. These challenges are exacerbated when most of the cohort is still living and must be censored in traditional survival analysis.

Here we overcome these issues by using pseudo restricted mean survival time (RMST) to identify genetic variants associated with resilience and longevity. Pseudo RMST applies a classical RMST framework for survival analysis while generating pseudo-observations for all animals, including animals that would otherwise be censored. This method is underutilized in the fields of aging and genetics despite addressing critical issues in the modeling of longevity in our best models of human resilience. To identify the genetic architecture underlying this resilience phenotype, we performed a whole-exome association analysis to identify high-impact changes contributing to longevity on all animals in the cohort.

## Methods

### Sample Description

The sample comprised 239 rhesus macaques (*Macaca mulatta*) from the Wake Forest Radiation Late Effects Cohort (RLEC) exposed to whole-body radiation in adolescence (Table 1). Irradiation occurred as part of a range of studies of mediators and treatments of acute radiation injury at multiple institutions. Following these acute studies, animals were donated to the RLEC, where they are maintained on a Western diet (Diet 5LOP, LabDiet) and monitored for long-term health effects (Olson et al., 2025). We have previously described the distribution of individual chronic diseases as well as multimorbidity in these animals (Quillen et al., 2025). Briefly, there are both increased rates and earlier onset of many common age-related, chronic conditions among the irradiated animals, including evidence of progression towards diabetes, heart disease, osteoporosis, arthritis, and cataracts. Additionally, irradiated animals show increased numbers of comorbid conditions spanning multiple organ systems. Despite this increased disease burden, lifespan for irradiated animals is highly variable and significantly overlaps lifespans observed in non-irradiated monkeys (Table 1). To generate and maintain this resource, all data were collected in accordance with the Animal Welfare Act and the U.S. Department of Health and Human Services “Guide for the Care and Use of Laboratory Animals” and approved by the Wake Forest University School of Medicine Institutional Animal Care and Use Committee.

**Table 1:**
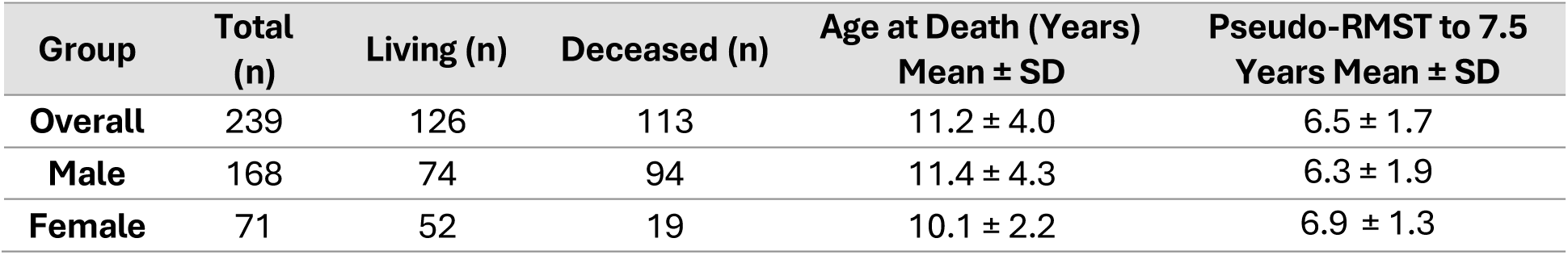
Pseudo-observation demographics.

| Group | Total (n) | Living (n) | Deceased (n) | Age at Death (Years) Mean ± SD | Pseudo-RMST to 7.5 Years Mean ± SD |
|---|---|---|---|---|---|
| **Overall** | 239 | 126 | 113 | 11.2 ± 4.0 | 6.5 ± 1.7 |
| **Male** | 168 | 74 | 94 | 11.4 ± 4.3 | 6.3 ± 1.9 |
| **Female** | 71 | 52 | 19 | 10.1 ± 2.2 | 6.9 ± 1.3 |

### Pseudo-Observation Generation

The basic concept behind the pseudo-observation (PO) approach is to overcome the challenge of censoring by creating a measurement of survival for every subject regardless of event or censoring status. This measurement is created through a transformation of the time-to-event data, taking the following form:

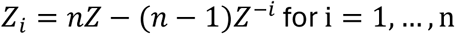

where *Z* is the global survival outcome of interest and *Z*^−*I*^ is the outcome of interest computed while excluding the *i^t^*^ℎ^ subject. A PO of this form can be computed for all subjects and can roughly be interpreted as the subject-specific contribution to whatever survival metric is being used. Then, this measurement can be utilized as an outcome variable in a generalized linear model (GLM) (Andersen, 2003).

Due to the challenges of estimating overall means in survival analysis in the presence of administrative right-censoring at study end, we use restricted mean survival time (RMST) and compute pseudo-RMST values as outcome variables. The term pseudo-mean (PM) will be used interchangeably with pseudo-RMST.

RMST, among other mean-based statistics, has become a recent fixture in survival analysis when considering how to circumvent the issues of modeling, testing, and interpreting survival data when hazards are expected to be non-proportional (NPH) (Magirr, 2021; Pak et al., 2017; Paukner C Chappell, 2021; Uno et al., 2014; Zhao et al., 2016). RMST can be computed and interpreted as the mean survival time from the beginning of follow-up to some time horizon *τ*. Thus, it estimates the expected survival duration of an individual until *τ*. This allows researchers to consider survival outcomes in terms of expected days, months, and years of life gained or lost based on relevant covariates. The purpose of using RMST rather than overall mean survival is to account for the limited follow-up time due to administrative right-censoring at study end. In time-to-event analysis, many subjects won’t have their event time recorded if they are either lost to follow-up (e.g. are sacrificed for non-clinical reasons) or the study ends prior to their event occurring. Thus, overall means cannot be directly estimated and may not be identifiable, though various restricted means can.

A restricted mean, 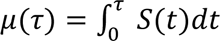, is graphically represented as the area under the survival curve from randomization *t* = 0 to some time horizon *t* = *τ*. In practice, *μ*(*τ*) can easily be estimated by computing the area under the Kaplan-Meier (KM) curve between 0 and *τ* as follows:

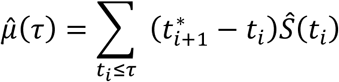

where *t_i_* is the *i^t^*^ℎ^ event time, 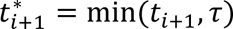 and 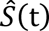 is the KM estimate of its respective survival function.

The subject-specific PMs to be used as the response variable *Z_i_*(*τ*) in our whole exome GLM analysis, where *τ* is the restriction time, is computed as:

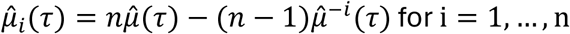

where 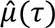 is the restricted mean computed based on the KM curve with all subjects and 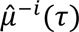 is the restricted mean computed based on the KM curve excluding subject *i* as in Equation 2. This particular technique was popularized by PK Andersen (Andersen et al., 2004; Andersen C Perme, 2010) and has been utilized by several others in dynamic risk prediction modeling in survival analysis (M. Lee et al., 2012; Paukner et al., 2024; Yang et al., 2021; Zhao et al., 2020). This PM can be interpreted as subject *i*’s individual contribution to the overall RMST.

### DNA Sequencing, Variant Calling, and Annotation

Whole-exome sequencing was performed at Baylor College of Medicine’s Human Genome Sequencing Center (BCM-HGSC) using one of two custom exome capture designs. The first combined NimbleGen SeqCap EZ HGSC VCRome 2.1 human exome probes with rhesus-specific spike-in probes (Caskey et al., 2019). The second combined Twist Bioscience HG38_HGSC_Twist_Comprehensive_Exome human exome probes with rhesus spike-in probes (Prall et al., 2024). Library preparation and target capture followed the manufacturers’ protocols, and captured libraries were sequenced on the Illumina NovaSeq platform. Analyses were restricted to regions targeted by both designs, defined as the intersection of the target region BED files for the two designs based on Mmul_10 coordinates.

The 150 bp paired-end reads were aligned to the Mmul_10 (rheMac10, GCF_003339765.1) rhesus genome assembly with BWA-MEM (v0.7.12), and duplicate reads were marked with Picard MarkDuplicates (v1.105). Single nucleotide variants (SNVs) and small insertion/deletions (indels) were called per sample with GATK HaplotypeCaller (v4.3.0.0) in GVCF mode. The resulting per-sample GVCFs were combined with

GenomicsDBImport and jointly genotyped with GenotypeGVCFs (McKenna, Hanna et al. 2010, DePristo, Banks et al. 2011). SNVs failing the GATK recommended hard filters (QD < 2.0, FS > 60.0, MQ < 40.0, MQRankSum < −12.5, ReadPosRankSum < −8.0, SOR > 3.0; https://gatk.broadinstitute.org/hc/en-us/articles/360035531112--How-to-Filter-variants-either-with-VQSR-or-by-hard-filtering) were removed. A total of 683,011 SNVs were retained for downstream analyses. Indels were not further examined. Raw reads are available through the NCBI Sequence Read Archive under BioProject accessions PRJNA1537671.

Variants were annotated with the Ensembl Variant Effect Predictor (VEP v108) using merged Ensembl and RefSeq models for Mmul_10 (McLaren et al., 2016). To assess variant effects in orthologous human genes, we projected variants onto the human genome (GRCh38) with Picard LiftoverVcf (v2.18.14) using the UCSC rheMac10ToHg38 chain file, with LIFTOVER_MIN_MATCH=0.95 and RECOVER_SWAPPED_REF_ALT=true. Liftover was performed reciprocally, and only sites that lifted back to their original rhesus position were retained. All SNVs that passed reciprocal liftover were annotated using Combined Annotation Dependent Depletion (CADD v1.6) (Kircher et al., 2014). Because of the extremely high homology between human and NHP protein-coding regions, we expect predicted changes in one species to cause similarly deleterious or benign changes in the other. This assumption is empirically supported by the finding that protein-altering variants shared between humans and non-human primates are benign in 99% of ClinVar cases; the functional impact of such variants is expected to be concordant across species (Gao et al., 2023; Sundaram et al., 2018).

### Whole-Exome Association Analysis

To identify genetic influences on longevity in our cohort, we analyzed pseudo-RMST based on survival to 7.5 years, as this provides the largest sample based on the age breakdown of living animals in the RLEC. We fit a linear mixed-effects model in GEMMA with the PM for each animal as the phenotype, sex as a fixed effect, and a standardized empirical relatedness matrix calculated based on all variants as a random effect (Zhou C Stephens, 2012). While the animals were assumed unrelated when acquired, empirical kinship estimates reflect genetic similarity among animals in our study equivalent to 5 pairs of first-degree, 36 second-degree, and 43 third-degree relatives. Associations were based on the likelihood ratio test using a genome-wide significance threshold of *p* < 1 × 10^-8.

To determine whether the significant variants represented independent genetic signals or were in linkage disequilibrium (LD), we performed a pairwise LD analysis using PLINK (v1.90b6.21) (Purcell et al., 2007). We calculated the pairwise squared correlation coefficient (R^2^) among all significant variants using a 50-kb window. Variants with an R^2^ > 0.8 were considered to be in strong LD. For the purpose of defining independent loci, variants falling within the same high-LD block were collapsed into a single locus. All other variants not meeting the LD threshold were considered independent singletons.

### Gene Set Enrichment

To predict the functional context of the single-variant association results at the pathway level, we performed a ranked Gene Set Enrichment Analysis (GSEA) using the clusterProfiler R package (v4.18.4) (Yu et al., 2012). We generated three complementary ranking metrics for each gene to ensure robustness of the enrichment findings and reduce false positives driven by a single metric (1) absolute effect size (∣ *β* ∣), (2) statistical significance (−log_10_(*p_LRT_*)), and (3) a combined score, defined as: ∣ *β* ∣× (−log_10_(*p_LRT_*)).

To obtain a single representative value per gene, we filtered out variants that were not mapped to the human genome and retained the variant with the lowest p-value from the GEMMA likelihood ratio test for each Ensembl ID (removing version suffixes) using dplyr (v1.2.1). The mapped Ensembl IDs were converted to Entrez IDs using the bitr function from clusterProfiler with the org.Hs.eg.db annotation database (v3.22.0) (Carlson, 2017) to align with the Kyoto Encyclopedia of Genes and Genomes (KEGG) human pathway database (hsa) (Ogata et al., 1999). Duplicated Entrez IDs were resolved by retaining the ID with the largest absolute beta value.

For each of the three ranking metrics, we constructed a ranked gene list and performed GSEA against the KEGG database using the gseKEGG function in clusterProfiler. We used 1,000 gene-set permutations and retained pathways with a false discovery rate (FDR, Benjamini-Hochberg) less than 0.05 as significantly enriched.

### Pathology and Fibrosis Scoring

To assess the physiological manifestations of genetically predicted fibrosis, we evaluated the presence and severity of pulmonary and myocardial fibrosis at time of death. This included 124 RLEC animals that died between 2008 and 2026, consisting of 100 irradiated and 24 non-irradiated control animals (Table 2).

**Table 2:** Fibrosis demographics.

| **Group** | **Sample Size** |  |  | **Mean Age at Death ± SD (Years)** |  |  |
|---|---|---|---|---|---|---|
|  | **Total (n)** | **Irradiated (n)** | **Non-Irradiated (n)** | **Total** | **Irradiated** | **Non-Irradiated** |
| **Overall** | 124 | 100 | 24 | 12.5 ± 4.6 | 11.5 ± 4.2 | 16.7 ± 4.0 |
| **Male** | 104 | 84 | 20 | 12.5 ± 4.8 | 11.6 ± 4.5 | 16.3 ± 4.2 |
| **Female** | 20 | 16 | 4 | 12.3 ± 4.2 | 10.7 ± 2.8 | 18.6 ± 2.3 |

Following euthanasia or spontaneous death, all animals underwent a comprehensive, standardized necropsy, as described in Schaaf et al., 2025 (Schaaf et al., 2025). A complete gross examination was performed, followed by systematic histologic evaluation of major organ systems, including the heart (right ventricle, left ventricle, and interventricular septum) and the lungs (one section from all six lobes). Necropsy reports and histologic slides for all animals were reviewed by a board-certified veterinary pathologist.

Fibrosis severity in the myocardial and pulmonary tissues was graded on a 4-point ordinal scale: 0 = none, 1 = mild/minimal, 2 = moderate, and 3 = severe/marked. When multiple grades were reported for a single tissue, the highest observed grade was recorded. To reduce inter-rater variability and ensure consistency, all scoring was performed by a single pathologist (GWS).

### Gene Burden and SKAT-O Analysis

Based on the results of the pathway analysis, we investigated the role of two enriched pathways in fibrosis severity: cadherin signaling (KEGG hsa04519) and integrin signaling (KEGG hsa04518). Analyses were performed on two cohorts: all animals, including irradiated and control, with available fibrosis scoring data (n = 124) and a subset of animals that received any radiation dose (dose > 0 Gy) (n = 100). Genotypes were extracted from filtered VCF files using *vcfR* (v1.16.0) (Knaus C Grünwald, 2017) and converted to numeric allele counts (0, 1, or 2), representing the number of alternative alleles carried by each animal at each variant site. Data manipulation, including joining variant annotations and phenotype data, was performed using *dplyr*(v1.2.1) and *tidyr* (v1.3.2) (Wickham et al., 2019).

For each pathway, analysis was restricted to its GSEA-derived leading-edge genes – the subset of pathway genes that drove the enrichment signal -to aggregate variant effects and capture multiple potentially rare but impactful variants that may collectively contribute to increased fibrosis susceptibility. We selected the leading-edge gene set for each pathway based on the GSEA metric under which it demonstrated the most robust enrichment. For the cadherin pathway, which was significantly enriched across multiple metrics, we evaluated leading-edge gene sets from both the absolute effect size and combined rankings as a sensitivity analysis. For the integrin pathway, which reached significance only under the statistical significance ranking, we utilized the leading-edge gene set derived from that metric.

Variants were weighted according to their CADD scores, allowing us to prioritize likely biologically relevant variants over benign ones. We compared three weighting schemes: (1) an unweighted model in which all variants were assigned a weight of 1, serving as a baseline for standard burden test; (2) a linear model in which variant’s weights were defined as the variant’s raw CADD score divided by the maximum raw CADD score across all variants of all genes in the dataset; and (3) a threshold model where variants with a CADD PHRED score greater than 20 were assigned a weight of 1, while all others were assigned 0. Selecting a threshold value of CADD PHRED scores greater than 20 retains the top 1% most deleterious variants (Kircher et al., 2014).

For total burden tests, a total pathway burden score was computed for each animal by summing the weighted allele counts across all leading-edge genes in the pathway. The primary phenotypic traits were myocardial fibrosis score and pulmonary fibrosis score, both measured on an ordinal scale from 0 to 3. As fibrosis severity and prevalence increase with age, all statistical models were adjusted for age at death to prevent confounding and isolate the genetic effects. Linear regression was used to test the association between total pathway-wide burden and each fibrosis phenotype.

Per-gene analyses were performed using Sequence Kernel Association Test-Optimal (SKAT-O v2.2.5) (S. Lee et al., 2012), an adaptive combination of the burden test and the variance-component SKAT that is sensitive to the direction of the variant effects within a gene. For each gene, we constructed a sample-by-variant genotype matrix and applied SKAT-O with the CADD-derived variant weights for the given scheme. A null model adjusting for age at death was fit for each cohort and phenotype combination, and each gene was then tested against that null model. The Benjamini-Hochberg method was applied within each combination of cohort, fibrosis phenotype, and weighting scheme. Genes with an FDR-adjusted p-value < 0.10 were considered nominally significant in this exploratory analysis. Visualizations were generated using *ggplot2* (v4.0.3) (Wickham, 2016) and *ggrepel* (v0.9.8) (Slowikowski, 2016). All analyses were performed in R (version 4.5.1) (R Core Team, 2025).

## Results

When analyses are limited to only animals with a known healthspan, we eliminate 126 animals that are still living or were euthanized for reasons unrelated to severe health decline. Whole exome analysis in this reduced cohort of 113 animals fails to identify a single variant with genome-wide significance. In contrast, pseudo-observations could be determined for all 239 irradiated animals, resulting in 63 autosomal single-nucleotide variants (SNVs) significantly associated with longevity (Figure 2).

Since whole-exome association analyses can identify multiple highly correlated variants within the same genomic region, we assessed linkage disequilibrium (LD) among the 63 significant autosomal SNVs to determine the number of independent genetic signals. LD identified 7 high-LD blocks that collapsed 20 SNVs. After collapsing these blocks into single loci, the 63 SNVs represent 50 unique autosomal genomic loci (7 LD blocks + 43 singletons). Of these 50 independent loci, 9 either failed reciprocal liftover or did not map to a human gene or open reading frame. The remaining 41 loci mapped to 35 unique human-equivalent genes or open reading frames, as several independent loci mapped to different regions of the same gene. Supplementary Table 1 lists significant longevity-associated variants with annotations.

**Figure 1.**
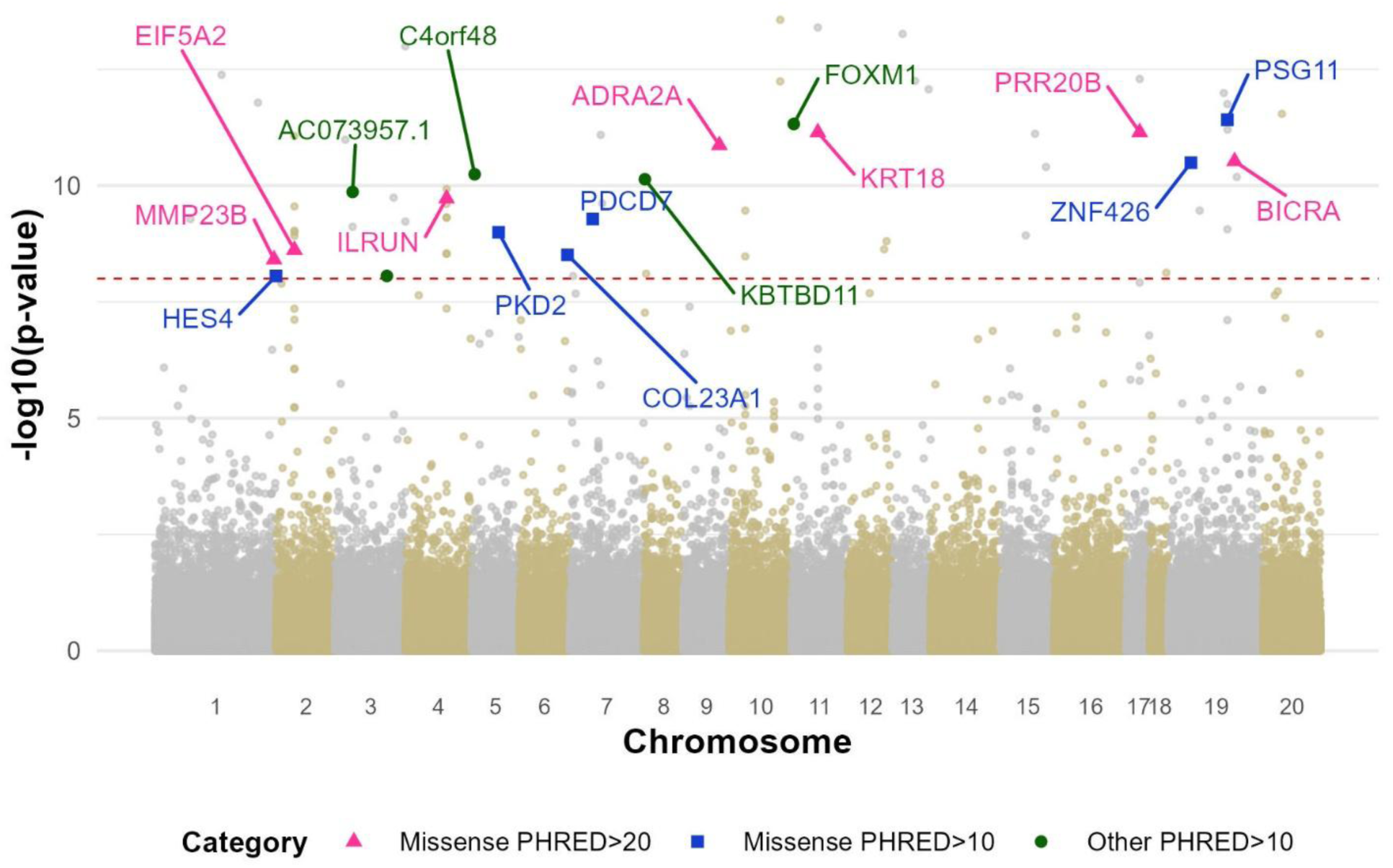
Manhattan plot of whole-exome associations with annotation of variants predicted to have the largest effect based on CADD PHRED scores.

**Table 2.**
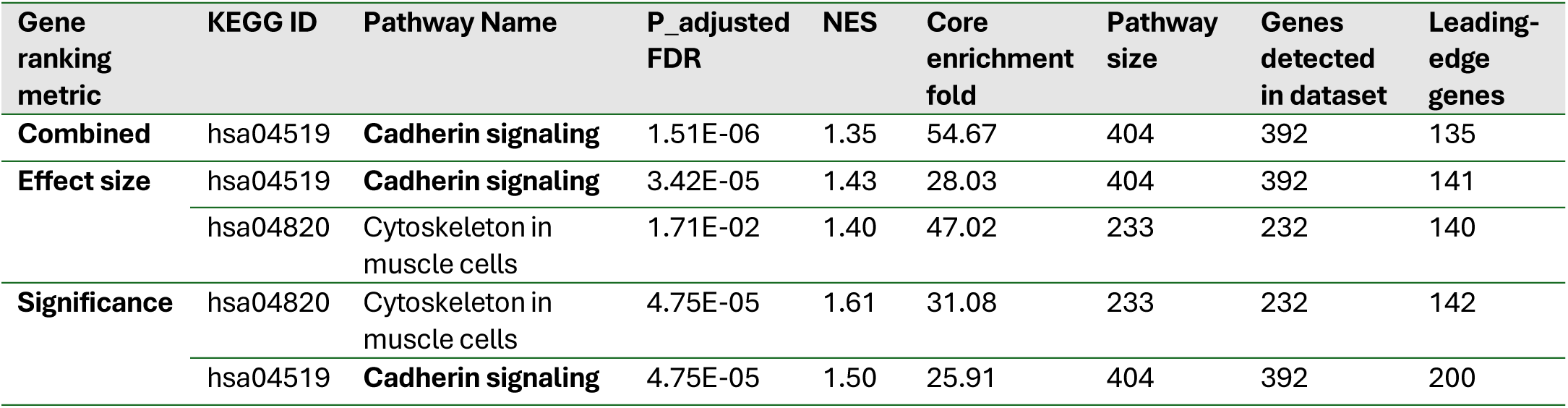
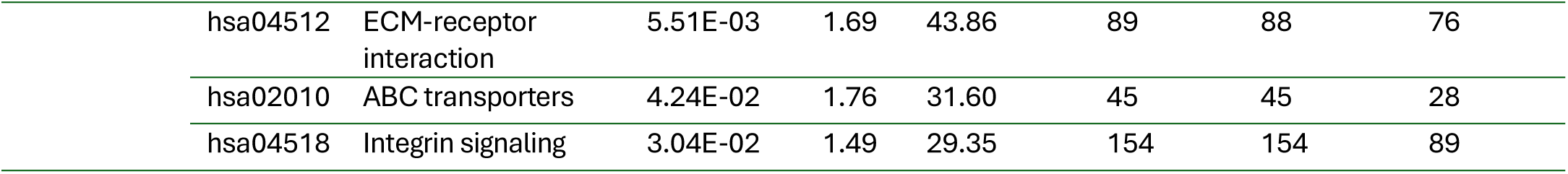
Summary of significantly enriched KEGG pathways (FDR < 0.05) from GSEA using three ranking metrics (effect size, significance, and combined), showing enrichment scores (NES), core enrichment fold, pathway sizes, and leading-edge gene counts.

| Gene ranking metric | KEGG ID | Pathway Name | P_adjusted FDR | NES | Core enrichment fold | Pathway size | Genes detected in dataset | Leading-edge genes |
|---|---|---|---|---|---|---|---|---|
| **Combined** | hsa04519 | **Cadherin signaling** | 1.51E-06 | 1.35 | 54.67 | 404 | 392 | 135 |
| **Effect size** | hsa04519 | **Cadherin signaling** | 3.42E-05 | 1.43 | 28.03 | 404 | 392 | 141 |
|  | hsa04820 | Cytoskeleton in muscle cells | 1.71E-02 | 1.40 | 47.02 | 233 | 232 | 140 |
| **Significance** | hsa04820 | Cytoskeleton in muscle cells | 4.75E-05 | 1.61 | 31.08 | 233 | 232 | 142 |
|  | hsa04519 | **Cadherin signaling** | 4.75E-05 | 1.50 | 25.91 | 404 | 392 | 200 |
|  | hsa04512 | ECM-receptor interaction | 5.51E-03 | 1.69 | 43.86 | 89 | 88 | 76 |
|  | hsa02010 | ABC transporters | 4.24E-02 | 1.76 | 31.60 | 45 | 45 | 28 |
|  | hsa04518 | Integrin signaling | 3.04E-02 | 1.49 | 29.35 | 154 | 154 | 89 |

To translate single-variant association signals into biologically interpretable pathways, we performed a Gene Set Enrichment Analysis (GSEA) on the gene-level summary statistics. The number of significantly enriched KEGG pathways (FDR < 0.05) varied across ranking metrics: statistical significance identified the greatest number (n = 5), followed by the absolute effect size (n = 2), and the combined score (n = 1) (Table 2).

We quantified pathway enrichment using the Normalized Enrichment Score (NES), where positive values indicate enrichment of pathway genes and larger magnitudes indicate stronger enrichment (Subramanian et al., 2005). The Cadherin signaling pathway (hsa04519) demonstrated robust cross-metric enrichment, reaching significance under all three ranking methods. This reproducibility across ranking strategies, combined with its strong enrichment signal, identified Cadherin signaling as the most robust pathway association.

The Cytoskeleton in muscle cells pathway (hsa04820) was significant under both the effect size and significance rankings. Under significance-based ranking, three additional pathways were enriched: ECM-receptor interaction (hsa04512), ABC transporters (hsa02010), and Integrin signaling (hsa04518).

Among the 124 animals with available fibrosis scores (described in Figure 2), 3,246 variants mapped to the absolute effect size GSEA leading-edge genes of the cadherin signaling pathway, representing 137 unique genes with at least one variant. For the integrin pathway, 3,936 variants mapped to 89 leading-edge genes, based on the statistical significance GSEA metric, with at least one variant present in the cohort. To test whether the accumulation of these variants influenced disease pathology, we evaluated fibrosis severity in these animals. Fibrosis scores ranged from 0 to 3 for both tissues, with higher scores observed more frequently in irradiated animals, particularly in the myocardial tissue. As shown in Figure 2, non-irradiated animals were generally older at death and exhibited a broader distribution of fibrosis scores, though the overall severity was lower than in that of irradiated animals.

**Figure 2:**
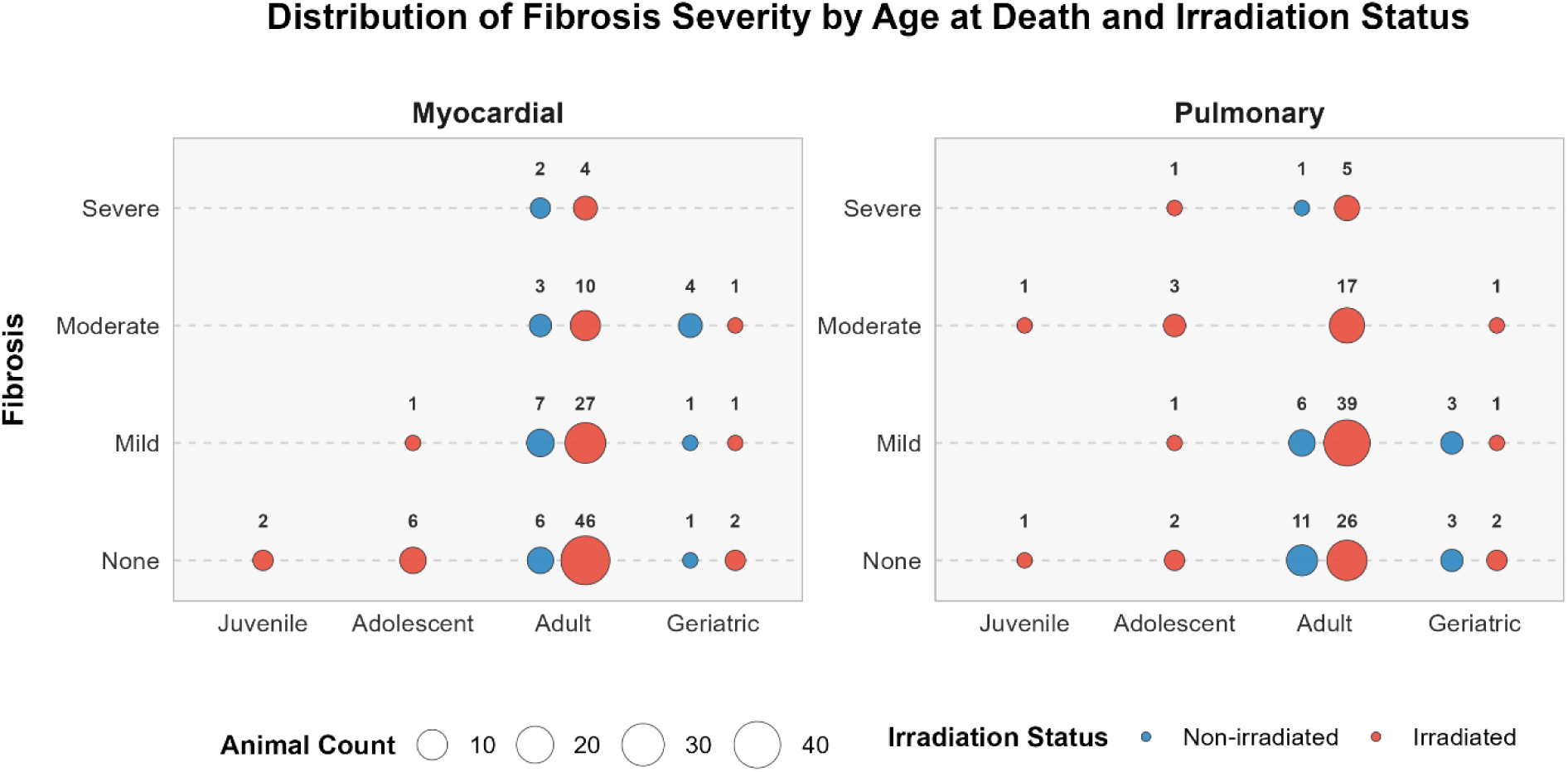
Distribution of fibrosis scores by age at death and irradiation status. Dot size reflects the number of animals per group; exact counts are shown above each dot. Age categories: Juvenile: ≤4 years; Adolescent: 4–7 years; Adult: 7–20 years; Geriatric: >20 years. Fibrosis was scored on a 4-point ordinal scale: 0 = none, 1 = mild/minimal, 2 = moderate, 3 = severe/marked (when multiple grades were reported, the highest grade was recorded). Non-irradiated animals are shown in blue; irradiated animals in red. Separate panels show myocardial and pulmonary fibrosis. Irradiated animals had higher fibrosis scores across age groups, particularly in myocardial tissue, with non-irradiated animals generally older at death.

Total burden tests assessed whether the cumulative CADD-weighted burden of variants across the leading-edge cadherin and integrin genes was associated with fibrosis severity (Table 3). In the cadherin pathway, total burden was significantly associated with pulmonary fibrosis in both the all-animal and irradiated-only cohorts under the threshold and linear schemes. For myocardial fibrosis in the cadherin pathway, significant associations were observed in the all-animal cohort under both the threshold and linear schemes, while the irradiated-only cohort showed no significant associations under any weighting scheme. In the integrin pathway, total burden was significantly associated with pulmonary fibrosis in the all-animal cohort under the threshold and linear schemes, and in the irradiated-only cohort under the threshold scheme. Conversely, no weighting scheme reached statistical significance for myocardial fibrosis in the integrin pathway across either cohort.

**Table 3.** Total Burden Results.

| **Pathway** | **Fibrosis Phenotype** | **Animal Group** | **Weighing Scheme** | **p-value** | **Beta** | **SE** | **R<sup>2</sup>** |
|---|---|---|---|---|---|---|---|
| **Cadherin** | Pulmonary | all | threshold | **0.0273** | 0.0181 | 0.0081 | 0.0503 |
|  |  |  | linear | **0.0422** | 0.0220 | 0.0107 | 0.0444 |
|  |  |  | unweighted | 0.0537 | 0.0017 | 0.0009 | 0.0412 |
|  |  | irradiated | threshold | **0.0262** | 0.0213 | 0.0094 | 0.0500 |
|  |  |  | linear | **0.0335** | 0.0272 | 0.0126 | 0.0457 |
|  |  |  | unweighted | 0.0941 | 0.0018 | 0.0011 | 0.0286 |
|  | Myocardial | all | threshold | **0.0356** | 0.0213 | 0.0100 | 0.1801 |
|  |  |  | linear | **0.0472** | 0.0153 | 0.0076 | 0.1768 |
|  |  |  | unweighted | 0.1533 | 0.0012 | 0.0012 | 0.1638 |
|  |  | irradiated | threshold | 0.1276 | 0.0132 | 0.0086 | 0.1254 |
|  |  |  | linear | 0.1302 | 0.0175 | 0.0115 | 0.1251 |
|  |  |  | unweighted | 0.3919 | 0.0008 | 0.0010 | 0.1109 |
| **Integrin** | Pulmonary | all | threshold | **0.0085** | 0.0270 | 0.0101 | 0.0663 |
|  |  |  | linear | **0.0321** | 0.0168 | 0.0077 | 0.0481 |
|  |  |  | unweighted | 0.0623 | 0.0013 | 0.0007 | 0.0392 |
|  |  | irradiated | threshold | **0.0203** | 0.0283 | 0.0120 | 0.0543 |
|  |  |  | linear | 0.0688 | 0.0015 | 0.0008 | 0.0337 |
|  |  |  | unweighted | 0.0910 | 0.0159 | 0.0093 | 0.0292 |
|  | Myocardial | all | threshold | 0.0543 | 0.0141 | 0.0073 | 0.1752 |
|  |  |  | linear | 0.0975 | 0.0161 | 0.0096 | 0.1686 |
|  |  |  | unweighted | 0.2006 | 0.0008 | 0.0007 | 0.1610 |
|  |  | irradiated | threshold | 0.2574 | 0.0096 | 0.2574 | 0.1160 |
|  |  |  | linear | 0.5258 | 0.0005 | 0.5258 | 0.1079 |
|  |  |  | unweighted | 0.5470 | 0.0067 | 0.0111 | 0.1075 |

As a sensitivity analysis, we also assessed total burden in the cadherin pathway using the combined-metric GSEA leading-edge gene set (Supplementary Table 2). These results generally mirrored the primary analysis, with total burden significantly associated with pulmonary fibrosis under the threshold scheme for the all-animal cohort and under both the threshold and linear schemes for the irradiated-only cohort, and with myocardial fibrosis in the all-animal cohort under the linear scheme.

Across both pathways, significant associations were consistently restricted to the threshold and linear weighting schemes, with the unweighted control scheme remaining non-significant in all comparisons. Total burden collapses all pathway genes into a single score and therefore cannot distinguish risk-conferring from protective contributions.

Per-gene SKAT-O analyses identified several genes significantly associated with fibrosis phenotypes across both the cadherin and integrin pathways (Figure 3 and Table 4). In the cadherin pathway, using the leading-edge genes derived from the absolute effect size GSEA ranking, *PCDHGA5* demonstrated a consistently positive association with myocardial fibrosis in the all-animal cohort, reaching significance under all three weighting schemes. *PCDHGB2* and *PSENEN* also showed positive associations with myocardial fibrosis under the unweighted and threshold schemes, respectively, indicating that a greater burden of variants in these genes correlates with increased fibrosis severity. Conversely, *PCDHA11* was negatively associated with myocardial fibrosis under the threshold scheme, suggesting a protective effect. As a sensitivity analysis, we also tested the cadherin leading-edge genes derived from the combined GSEA ranking. In this gene set, PCDHGA3 and PSENEN were positively associated with myocardial fibrosis in the all-animal cohort under the threshold weighting scheme.

**Figure 3:**
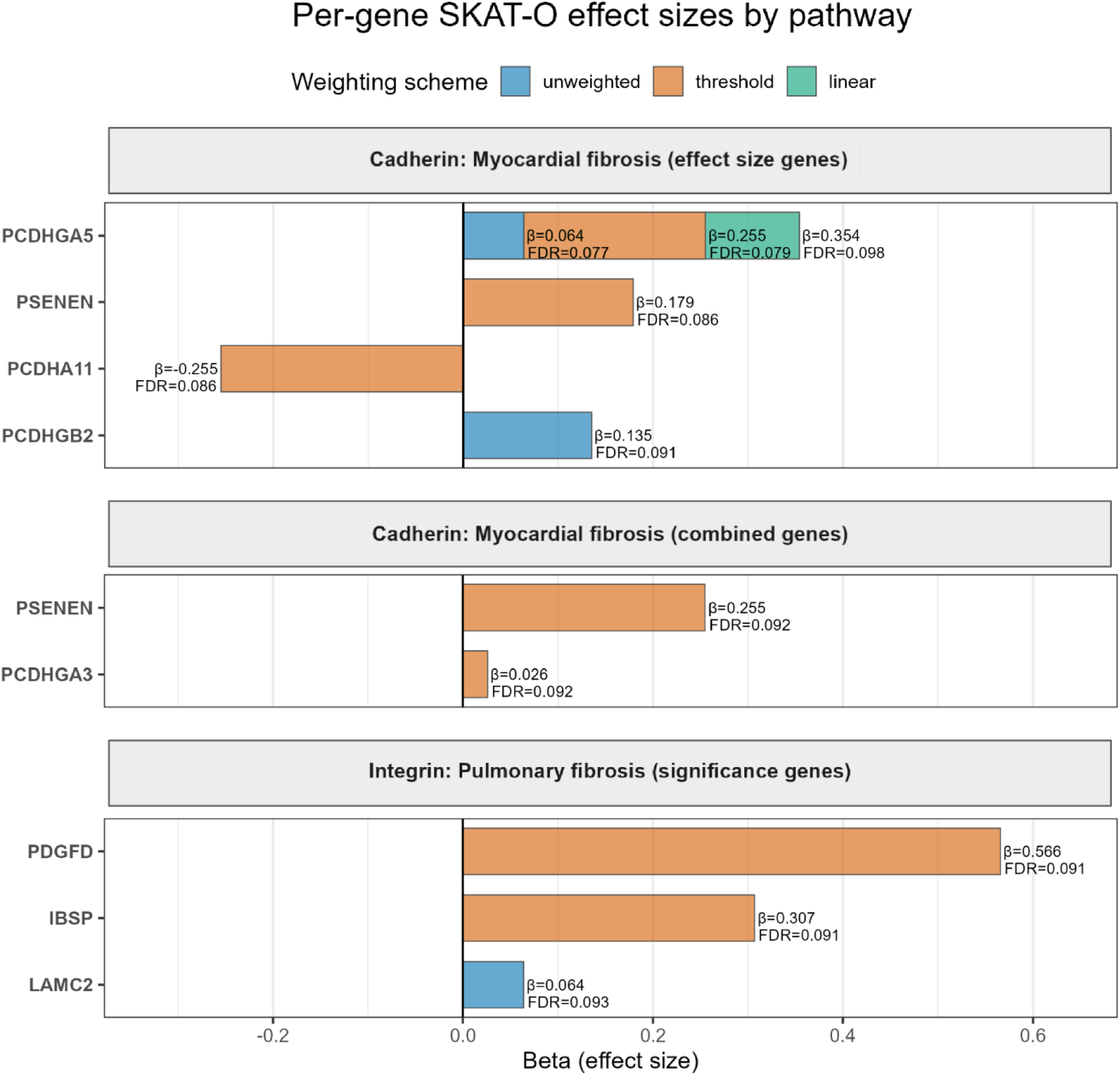
Per-gene SKAT-O effect sizes by pathway. Significant gene-level associations (FDR < 0.10) with fibrosis severity are shown for the GSEA leading-edge genes. The top panel shows myocardial fibrosis associations for the cadherin pathway using the absolute effect size metric leading-edge genes; the middle panel shows myocardial fibrosis associations for the cadherin pathway using the combined-metric leading-edge genes (sensitivity analysis); the bottom panel shows pulmonary fibrosis associations for the integrin pathway using the significance-metric leading-edge genes. Bars represent the beta coefficient (effect size) derived from the burden regression, oriented to show the direction of effect (positive values indicate greater variant burden is associated with more severe fibrosis; negative values indicate a protective association). Bar color indicates the weighting scheme under which the gene reached significance (blue = unweighted, orange = threshold, green = linear). Exact beta coefficients and FDR-adjusted p-values are annotated for each association.

**Table 4.** Per-gene SKAT-O Results.

| Pathway | Phenotype | Cohort | GSEA Method | Gene | Beta | p-value | P adj FDR | Carriers (n) | Weighting Scheme |
|---|---|---|---|---|---|---|---|---|---|
| Cadherin | Myocardial fibrosis | all | Effect size | **PCDHGA5** | 0.0637 | 0.0006 | **0.0771** | 116 | unweighted |
|  |  |  |  |  | 0.2550 | 0.0008 | **0.0792** | 82 | threshold |
|  |  |  |  |  | 0.3540 | 0.0007 | **0.0983** | 111 | linear |
|  |  |  |  | PCDHA11 | -0.2550 | 0.0022 | **0.0859** | 88 | threshold |
|  |  |  |  | PCDHGB2 | 0.1350 | 0.0013 | **0.0908** | 124 | unweighted |
|  |  |  |  | **PSENEN** | 0.1790 | 0.0025 | **0.0859** | 65 | threshold |
|  |  |  | Combined |  | 0.255 | 0.0025 | **0.0922** | 65 | threshold |
|  |  |  |  | PCDHGA3 | 0.0258 | 0.002 | **0.0922** | 124 | threshold |
| Integrin | Pulmonary fibrosis | all | Significance | IBSP | 0.3070 | 0.0021 | **0.0906** | 40 | threshold |
|  |  |  |  | PDGFD | 0.5660 | 0.0024 | **0.0906** | 6 | threshold |
|  |  |  |  | LAMC2 | 0.0639 | 0.0011 | **0.0933** | 123 | unweighted |

In the integrin pathway, per-gene SKAT-O identified three genes positively associated with pulmonary fibrosis in the all-animal cohort using the significance GSEA ranking-derived leading-edge gene set: *IBSP* and *PDGFD* under the threshold scheme, and *LAMC2* under the unweighted scheme. Notably, no per-gene associations reached significance for pulmonary fibrosis in the cadherin pathway, for myocardial fibrosis in the integrin pathway, or in the irradiated-only cohort for any phenotype. All associations were adjusted for age at death, confirming that the observed genetic effects are independent of age-related confounding.

## Discussion

This study establishes pseudo-RMST as a valuable tool for genetic association studies in aging cohorts. Although pseudo-observations have been employed in survival analysis for clinical epidemiology (e.g. (Choi et al., 2022)), their potential for human or non-human primate genetic analyses has, to our knowledge, remained unexplored. Application of pseudo-RMST to this non-human primate model generated pseudo-observations for all animals, including those whose event times were administratively censored, allowing retention of the full cohort in our genetic analyses and thus overcoming the statistical power limitations inherent to traditional survival approaches. Indeed, analysis restricted to uncensored animals yielded no significant associations, whereas pseudo-RMST enabled the detection of variants contributing to resilience. Furthermore, restricting analyses to uncensored animals is likely to produce biased results, as censored and uncensored animals are likely to have different underlying genetic risks. This methodological framework forms the foundation for all subsequent biological discoveries.

The pseudo-RMST association analysis identified 63 autosomal variants significantly associated with longevity. An LD analysis confirmed that these variants represent 50 independent genomic loci, mapping to 35 unique human-equivalent genes. The lack of cross-locus LD among these hits reinforces that the genetic architecture of resilience in this model is highly polygenic. The associated variants predicted to be deleterious cluster in several genes previously linked to aging, including *ILRUN, EIF5A2, KRT18, BICRA, PRR20B, MMP23B, ADRA2A, PKD2, COL23A1, C4orf48, PDCD7, FOXM1, KBTBD11, ESYT2, NPIPA3, PSG4,* and *APOL1*. The presence of multiple independent loci within some of these genes (such as ILRUN and EIF5A2) further highlights their importance, as it suggests these regions harbor multiple functional variants independently contributing to the phenotype. However, the role of many others in promoting resilience and longevity was not previously investigated.

Of the 50 independent loci identified, 9 either failed reciprocal liftover or did not map to a known human gene or open reading frame. These unmapped loci may represent rhesus-specific regulatory regions, structural variants, or highly divergent sequences. While they could not be directly interpreted using human annotations, they represent potentially novel, primate-specific genetic contributors to resilience that warrant future investigation.

Our GSEA converged on five significantly enriched KEGG pathways: ECM-receptor interaction, cytoskeleton in muscle cells, ABC transporters, cadherin signaling, and integrin signaling. The ECM-receptor interaction pathway regulates the extracellular matrix (ECM), which provides structural support to tissues and serves as a dynamic signaling platform that regulates cell behavior through receptor-mediated interactions (De Luca, 2019). Age-related changes in the ECM, such as increased stiffness, can induce cellular senescence and contribute to the development of pathological conditions across organ systems (Guvatova et al., 2022). These include fibrotic and inflammatory disorders in the lung, liver, kidney, and heart, as well as cardiovascular disease, sarcopenia and osteoarthritis, skin disorders, neurodegenerative diseases, and cancer progression and metastasis (Guvatova et al., 2022). Among our candidate genes, *COL23A1* encodes a transmembrane collagen that directly participates in ECM-receptor interactions and has been associated with longevity, suggesting that structural integrity of the matrix is a determinant of resilience (Jabalameli et al., 2024). Furthermore, *MMP23B* (matrix metallopeptidase 23B) is involved in ECM degradation and its downregulation has been shown to inhibit the migration in cancer models (N. Li et al., 2024).

The cytoskeleton in muscle cells pathway provides the mechanical infrastructure that maintains cellular shape, enables intracellular transport, and mediates force transmission (Discher et al., 2005). During aging, altered actin expression disrupts cytoskeletal dynamics and mechanical properties of the cell surface, compromising the structural integrity of muscle fibers, which in turn impairs force transmission (Hughes et al., 2015; Kim et al., 2022). Several of our candidate genes fall within this pathway, linking them to loss of structural integration. *APOL1* variants have been linked to harmful actomyosin reorganization in podocytes (Pays, 2020), while *EIF5A2* depletion disrupts actin cytoskeletal organization and increases cell motility (Martínez-Férriz et al., 2023). *KRT18* encodes Keratin 18, which forms the structural keratin intermediate filament network involved in force transduction and organization (Jacob et al., 2018), and its levels are known to decline in aged muscle (Ganjayi et al., 2024). The enrichment of cytoskeletal pathway genes underscores the importance of maintaining mechanical homeostasis and structural integrity as a mechanism of resilience against age-related muscle dysfunction.

The ABC transporters pathway comprises ATP-binding cassette (ABC) transporters – membrane proteins that actively transport a wide range of substrates across cellular membranes, playing critical roles in cellular detoxification, maintaining lipid homeostasis, and protection of sensitive organs such as the brain (Efferth, 2003; Wada et al., 2025). With advancing age, cumulative exposure to environmental toxins and metabolic byproducts makes ABC transporter function increasingly critical. Within this pathway, our most significant longevity-associated gene was *ABCG1*. *ABCG1* is a key regulator of cholesterol efflux, and its deficiency in human macrophages promotes proinflammatory M1 polarization and activates the NLRP3 inflammasome (Kotlyarov C Kotlyarova, 2022). This directly links ABC transporter dysfunction to chronic inflammation, a core hallmark of aging (Baechle et al., 2023). Although fewer of our candidate genes mapped directly to this pathway, its enrichment alongside adhesion and structural pathways suggests that cellular housekeeping and detoxification work together with tissue integrity to combat the cumulative stresses of aging.

The cadherin signaling pathway demonstrated the most robust cross-metric enrichment in our analysis, and together with the integrin pathway, highlights the central role of cell adhesion in longevity. Cadherins are calcium-dependent adhesion molecules critical for tissue morphogenesis, maintenance of tissue integrity, and wound healing (Sivasankar C Xie, 2023). In aging, cadherin expression and function decline, leading to compromised tissue integrity and increased susceptibility to age-related diseases (Frantz et al., 2010). To understand how our longevity-associated genes might drive pathology, it is helpful to distinguish between cadherin subtypes. E-cadherin is the primary adhesion molecule in epithelial cells, where it forms the adherens junctions that maintain the epithelial barrier (Hu et al., 2023). Its dysregulation is a hallmark of epithelial-mesenchymal transition (EMT) and is strongly implicated in pulmonary fibrosis, where alveolar epithelial cells are a primary source of fibrotic remodeling (Hu et al., 2023). In contrast, the heart lacks epithelial cells; instead, cadherin-based adhesion in the myocardium is mediated primarily by N-cadherin at cardiomyocyte intercalated discs (J. Li et al., 2006), while endothelial integrity is maintained by VE-cadherin at endothelial adherens junctions (Lampugnani et al., 2017). While our longevity hits did not map to the *CDH1* (E-cadherin) or *CDH2* (N-cadherin) genes directly, they did map to genes that regulate cadherin function and processing. *KRT18* directly binds to and stabilizes cell-surface E-cadherin (Liang et al., 2023). Other candidate genes alter cadherin expression: *FOXM1* and *PKD2* overexpression are negatively correlated with E-cadherin expression (Meng et al., 2015; Z. Pang et al., 2019), *ILRUN* downregulates E-cadherin (Zhang et al., 2015), *PDCD7* knockout reduces E-cadherin expression (Peng et al., 2018), and *EIF5A2* knockdown upregulates E-cadherin (Zheng et al., 2021). Interestingly, *EIF5A2* has been implicated in aging through several studies suggesting *EIF5A2* could function as a transcriptional regulator of aging (Liu et al., 2023). Overexpression of *EIF5A2* in transgenic mice exhibited accelerated organismal aging (Liu et al., 2023). Additionally, *ILRUN*, a regulator of lipid metabolism and inflammation, contains causal variants that have been linked to frailty across different tissues (Mak et al., 2025).

The integrin signaling pathway regulates another cell adhesion molecule family. Integrins are transmembrane receptors that mediate cell-cell and cell-ECM adhesion, enabling bidirectional signal transduction that regulates cell survival, proliferation, and migration (X. Pang et al., 2023). Among our top gene hits, *APOL1* interacts with integrin signaling pathways through the Apolipoprotein L1-suPAR-Integrin Axis, which forms a tripartite complex that drives rapid progression of chronic kidney disease in kidney podocytes (Hayek et al., 2017).

Collectively, these five pathways converge on a biological theme: the maintenance of tissue architecture through cell adhesion, extracellular matrix organization, and cytoskeletal integrity. Cadherins and integrins provide adhesive connections, the ECM provides the structural scaffold, the cytoskeleton provides the mechanical infrastructure, and ABC transporters regulate the cellular microenvironment. This convergence suggests that genetic variation governing the structural fidelity of tissues may represent a core mechanism distinguishing resilient from vulnerable individuals in the context of radiation and aging stress. Because the loss of this structural integrity is the defining pathological feature of fibrosis (Antar et al., 2023), we hypothesized that damaging variants in the cadherin and integrin pathways would predispose animals to more severe fibrotic pathology (Agarwal, 2014; Hu et al., 2023).

To test whether the pathway-level enrichment translated into a physiological phenotype, we first examined whether the cumulative CADD-weighted burden of all variants across the leading-edge genes of each pathway was associated with fibrosis severity, adjusting for age at death. This total-burden approach aggregates weak but functionally related effects across the pathway, thereby increasing power to detect a coordinated genotype-phenotype signal. The cadherin pathway’s total burden was significantly associated with both pulmonary and myocardial fibrosis, while the integrin pathway’s total burden was associated with pulmonary fibrosis but not myocardial fibrosis. Importantly, because total burden collapses all pathway genes into a single score, it cannot distinguish risk-conferring from protective contributions. The significant total-burden associations therefore provide pathway-level validation for the GSEA signals while motivating gene-level analyses to identify the specific drivers and directions of effect.

Per-gene SKAT-O analyses revealed distinct genetic architectures for myocardial and pulmonary fibrosis. Within the cadherin signaling pathway, five genes reached our exploratory FDR threshold (FDR < 0.10) as significantly associated with myocardial fibrosis severity in the cadherin signaling pathway: *PSENEN*, *PCDHGA5*, *PCDHGB2*, *PCDHGA3*, and *PCDHA11*.

*PSENEN* was significantly and positively associated with myocardial fibrosis severity in the all-animal cohort under the threshold weighting scheme for both leading-edge gene sets (effect-size based and combined effect-size/significance based), underscoring its robustness. *PSENEN* (presenilin enhancer, gamma-secretase subunit) encodes the PEN-2 subunit of the gamma-secretase complex, whose functions include modulating Notch signaling (Serneels et al., 2024). Notch signaling is involved in the expression of pro-fibrotic markers and has been implicated in pulmonary, renal, hepatic, and cutaneous fibrosis (Condorelli et al., 2021). Additionally, the gamma-secretase complex plays an indispensable role in protocadherin processing (Pancho et al., 2020). Specifically, gamma-secretase cleaves the protocadherin protein, releasing a soluble protocadherin intracellular domain (ICD) that can translocate into the nucleus and regulate gene expression (W. V. Chen C Maniatis, 2013; Pancho et al., 2020).

The protocadherin gamma (PCDHG) cluster demonstrated a strong positive association with myocardial fibrosis in our cohort. *PCDHGA5* was robustly associated across all weighting schemes, while *PCDHGB2* (unweighted) and *PCDHGA3* (threshold -combined-metric leading-edge set) also reached statistical significance. Clustered protocadherins are calcium-dependent cell adhesion molecules classically studied in the nervous system for establishing neuronal connectivity (Flaherty C Maniatis, 2020; Pancho et al., 2020).

More recently however, gamma-protocadherins have been found to be expressed in cardiac fibroblasts, and their dysregulation is increasingly recognized in cardiac disease and inflammation (Chaudhari et al., 2025; Sandstedt et al., 2021). The positive associations observed suggest that damaging variants in these genes impair normal fibroblast cell-cell adhesion and intracellular signaling, predisposing the myocardium to increased fibrosis incidence following chronic stress and tissue injury (Chaudhari et al., 2025).

In contrast, *PCDHA11* was negatively associated with myocardial fibrosis severity under the threshold scheme. This negative association suggests a protective effect where increased burden of variants in this gene is associated with reduced myocardial fibrosis severity. *PCDHA11* (protocadherin alpha 11) encodes a member of the clustered alpha-protocadherins, which physically and functionally cooperate with gamma-protocadherins to regulate cell survival, adhesion, and contact-dependent signaling (Ing-Esteves et al., 2018; Pancho et al., 2020). Because these alpha-protocadherins directly participate in combinatorial complexes with gamma-protocadherins, damaging variants in *PCDHA11* may disrupt these surface interactions and decouple the pro-fibrotic signaling otherwise driven by the gamma-cluster.

In contrast to the myocardial associations observed in the cadherin pathway, per-gene SKAT-O analyses within the integrin pathway identified three genes positively associated with pulmonary fibrosis severity in the all-animal cohort: *IBSP*, *PDGFD*, and *LAMC2*. Unlike the cadherin genes, which primarily mediate cell-cell interactions, these three genes converge on cell-extracellular matrix interactions and growth factor signaling.

*IBSP* (integrin-binding sialoprotein) was positively associated with pulmonary fibrosis severity under the threshold scheme. *IBSP* encodes bone sialoprotein, an extracellular matrix glycoprotein that binds cell-surface integrins and collagen (Y. Chen et al., 2024). While traditionally studied in bone, recent proteome-wide Mendelian randomization by Chen et al. (2024) (Y. Chen et al., 2024) identified circulating IBSP as causally protective against idiopathic pulmonary fibrosis. In that study, experimental bleomycin-induced lung fibrosis caused a significant decrease in pulmonary *IBSP* expression, and in previous studies, *IBSP* knockout mice demonstrated elevated mRNA levels of *Col1a1*, a marker of fibrotic collagen deposition (Chavez et al., 2023; Y. Chen et al., 2024). This is consistent with a loss-of-function effect, where genetic disruption of IBSP impairs normal matrix organization and tissue homeostasis, worsening fibrotic remodeling after injury (Y. Chen et al., 2024).

*PDGFD* exhibited the largest effect size among the integrin-associated candidates under the threshold scheme. *PDGFD* encodes platelet-derived growth factor D, a growth factor that selectively binds and activates PDGFR-beta receptors, driving fibroblast proliferation, migration, and matrix deposition across various organs (Klinkhammer et al., 2018). In experimental models of lung fibrosis, *PDGFD* expression behaved unusually compared to other PDGF growth factor isoforms and became downregulated in the lung after injury (Charni Chaabane et al., 2014; Zhuo et al., 2004). This suppression is driven by TGF-beta1 and other pro-fibrotic cytokines, which act as negative feedback to restrain ongoing fibroblast proliferation (Charni Chaabane et al., 2014). Furthermore, blocking PDGFR-beta with inhibitors attenuates fibrocyte recruitment and reduces lung fibrosis in both bleomycin and radiation induced models (Klinkhammer et al., 2018). Therefore, damaging variants in *PDGFD* may disrupt this feedback circuit or alter receptor regulation; such variants could leave proliferation and scar formation unchecked, predisposing to more severe fibrosis.

*LAMC2* (laminin subunit gamma-2) was positively associated with pulmonary fibrosis severity under the unweighted model. *LAMC2* encodes the gamma2 chain of laminin-332, a specialized basement membrane protein critical for epithelial cell adhesion and tissue repair (Wang et al., 2026). In human lung disease, *LAMC2* is strongly expressed in regenerating alveolar epithelial cells in idiopathic pulmonary fibrosis (Wang et al., 2026). In a recent study, LAMC2 was shown to be upregulated in airway epithelial-mesenchymal transition (EMT) and collagen deposition through activation of the AKT pathway (Wang et al., 2026). In our cohort, the positive association suggests that damaging variants in *LAMC2* may alter baseline basement membrane stability or prime epithelial cells for EMT and matrix remodeling under chronic pulmonary stress.

Notably, no per-gene associations reached significance in the irradiated-only cohort for either pathway. This is likely due to reduced statistical power (n = 100) and the overwhelming effect of radiation on fibrosis severity. Irradiated animals exhibit significantly higher baseline fibrosis scores than non-irradiated controls at an earlier age, which may be affecting the ability to detect subtler genetic effects. In contrast, the full cohort of 124 animals includes non-irradiated controls with lower fibrosis scores that occur later in life, providing the variance necessary to detect genotype-phenotype associations.

Several limitations warrant consideration when interpreting these findings. First, while whole-exome sequencing captures protein-coding variation, the most likely region to harbor functional variants, we may have missed important regulatory variants in non-coding regions. Second, our fibrosis scores were assessed on an ordinal scale, which may not fully capture the continuous spectrum of fibrotic pathology. However, consistent scoring by a single pathologist reduces inter-rater variability and increases confidence in the ordinal assessments. Third, and importantly, the gene burden analysis was limited to the 124 deceased animals with fibrosis scoring data. This reduced sample size likely limited statistical power for detecting individual gene associations, potentially explaining why only a handful of genes reached our exploratory FDR threshold (0.10) and why none survived in the irradiated-only subset. Yet this limitation itself highlights the value of the pseudo-RMST approach used for the primary association analysis, which allowed retention of the full cohort of 239 animals and detection of the robust pathway-level signal that guided our subsequent validation.

Taken together, the total burden and per-gene results provide a genotype-to-phenotype bridge between the GSEA enrichment and the fibrotic pathology observed in this cohort. They suggest that variants in both cadherin and integrin pathway genes contributed to fibrosis severity in irradiated and non-irradiated animals, with accumulated variants exerting pro-fibrotic or protective effects. These per-gene findings are exploratory (FDR < 0.10) and require replication, but they offer a shortlist of candidate genes for future studies. Critically, we note that the genetic variants are not caused by the radiation but rather create an underlying risk for fibrosis that is then exacerbated by the acute and persistent stressors challenging these animals’ intrinsic resilience. Such stressors could involve radiation or chemotherapeutic treatments, smoking, or other environmental toxins and individual risk patterns could be critical in, for example, personalizing cancer treatments based on individual genetic risk of fibrosis. The association of longevity with genetic variants involved in fibrosis specifically and maintenance of tissue integrity broadly points to the importance of structural integrity of the cell in promoting longevity. Future research should expand beyond fibrosis to investigate other mechanisms by which these cellular maintenance genes support longevity.

This research was funded by NIH U01 AI150578, U19 AI67798, and R01 AG087957.

## References

Agarwal, S. K. (2014). Integrins and cadherins as therapeutic targets in fibrosis. Frontiers in Pharmacology, 5, 131. 10.3389/fphar.2014.00131

Andersen, P. K. (2003). Generalised linear models for correlated pseudo-observations, with applications to multi-state models. Biometrika, 90(1), 15–27. 10.1093/biomet/90.1.15

Andersen, P. K., Hansen, M. G., & Klein, J. P. (2004). Regression analysis of restricted mean survival time based on pseudo-observations. Lifetime Data Analysis, 10(4), 335–350. 10.1007/s10985-004-4771-0

Andersen, P. K., & Perme, M. P. (2010). Pseudo-observations in survival analysis. Statistical Methods in Medical Research, 19(1), 71–99. 10.1177/0962280209105020

Antar, S. A., Ashour, N. A., Marawan, M. E., & Al-Karmalawy, A. A. (2023). Fibrosis: Types, Effects, Markers, Mechanisms for Disease Progression, and Its Relation with Oxidative Stress, Immunity, and Inflammation. International Journal of Molecular Sciences, 24(4), 4004. 10.3390/ijms24044004

Baechle, J. J., Chen, N., Makhijani, P., Winer, S., Furman, D., & Winer, D. A. (2023). Chronic inflammation and the hallmarks of aging. Molecular Metabolism, 74, 101755. 10.1016/j.molmet.2023.101755

Carlson, M. (2017). Org.Hs.eg.db [Computer software]. Bioconductor. 10.18129/B9.BIOC.ORG.HS.EG.DB

Caskey, J. R., Wiseman, R. W., Karl, J. A., Baker, D. A., Lee, T., Maddox, R. J., Raveendran, M., Harris, R. A., Hu, J., Muzny, D. M., Rogers, J., & O’Connor, D. H. (2019). MHC genotyping from rhesus macaque exome sequences. Immunogenetics, 71(8–9), 531–544. 10.1007/s00251-019-01125-w

Charni Chaabane, S., Coomans de Brachène, A., Essaghir, A., Velghe, A., Lo Re, S., Stockis, J., Lucas, S., Khachigian, L. M., Huaux, F., & Demoulin, J.-B. (2014). PDGF-D expression is down-regulated by TGFβ in fibroblasts. PloS One, 9(10), e108656. 10.1371/journal.pone.0108656

Chaudhari, A., Axelsson, C., Mattsson Hultén, L., & Rotter Sopasakis, V. (2025). Toll-like Receptors 1, 3 and 7 Activate Distinct Genetic Features of NF-κB Signaling and γ-Protocadherin Expression in Human Cardiac Fibroblasts. Inffammation, 48(5), 2896–2907. 10.1007/s10753-025-02238-z

Chavez, M. B., Tan, M. H., Kolli, T. N., Zachariadou, C., Farah, F., Mohamed, F. F., Chu, E. Y., & Foster, B. L. (2023). Bone Sialoprotein Is Critical for Alveolar Bone Healing in Mice. Journal of Dental Research, 102(2), 187–196. 10.1177/00220345221126716

Chen, W. V., & Maniatis, T. (2013). Clustered protocadherins. Development, 140(16), 3297–3302. 10.1242/dev.090621

Chen, Y., Cao, S., Shao, S., & Tong, Z. (2024). Identifying prothrombin and bone sialoprotein as potential drug targets for idiopathic pulmonary fibrosis. BMC Pulmonary Medicine, 24(1), 488. 10.1186/s12890-024-03289-y

Choi, S., Choi, T., Lee, H.-Y., Han, S. W., & Bandyopadhyay, D. (2022). Doubly-robust methods for differences in restricted mean lifetimes using pseudo-observations. Pharmaceutical Statistics, 21(6), 1185–1198. 10.1002/pst.2223

Condorelli, A. G., El Hachem, M., Zambruno, G., Nystrom, A., Candi, E., & Castiglia, D. (2021). Notch-ing up knowledge on molecular mechanisms of skin fibrosis: Focus on the multifaceted Notch signalling pathway. Journal of Biomedical Science, 28(1), 36. 10.1186/s12929-021-00732-8

De Luca, M. (2019). The role of the cell-matrix interface in aging and its interaction with the renin-angiotensin system in the aged vasculature. Mechanisms of Ageing and Development, 177, 66–73. 10.1016/j.mad.2018.04.002

Discher, D. E., Janmey, P., & Wang, Y.-L. (2005). Tissue cells feel and respond to the stiffness of their substrate. Science, 310(5751), 1139–1143. 10.1126/science.1116995

Efferth, T. (2003). Adenosine triphosphate-binding cassette transporter genes in ageing and age-related diseases. Ageing Research Reviews, 2(1), 11–24. 10.1016/s1568-1637(02)00046-6

Flaherty, E., & Maniatis, T. (2020). The role of clustered protocadherins in neurodevelopment and neuropsychiatric diseases. Current Opinion in Genetics & Development, 65, 144–150. 10.1016/j.gde.2020.05.041

Frantz, C., Stewart, K. M., & Weaver, V. M. (2010). The extracellular matrix at a glance. Journal of Cell Science, 123(Pt 24), 4195–4200. 10.1242/jcs.023820

Ganjayi, M. S., Frank, S. W., Krauss, T. A., York, M. L., Bloch, R. J., & Baumann, C. W. (2024). Skeletal muscle adaptations following eccentric contractions are not mediated by keratin 18. Journal of Applied Physiology, 137(4), 903–909. 10.1152/japplphysiol.00496.2024

Gao, H., Hamp, T., Ede, J., Schraiber, J. G., McRae, J., Singer-Berk, M., Yang, Y., Dietrich, A. S. D., Fiziev, P. P., Kuderna, L. F. K., Sundaram, L., Wu, Y., Adhikari, A., Field, Y., Chen, C., Batzoglou, S., Aguet, F., Lemire, G., Reimers, R., … Farh, K. K.-H. (2023). The landscape of tolerated genetic variation in humans and primates. Science, 380(6648), eabn8153. 10.1126/science.abn8197

Guvatova, Z. G., Borisov, P. V., Alekseev, A. A., & Moskalev, A. A. (2022). Age-Related Changes in Extracellular Matrix. Biochemistry. Biokhimiia, 87(12), 1535–1551. 10.1134/S0006297922120112

Hayek, S. S., Koh, K. H., Grams, M. E., Wei, C., Ko, Y.-A., Li, J., Samelko, B., Lee, H., Dande, R. R., Lee, H. W., Hahm, E., Peev, V., Tracy, M., Tardi, N. J., Gupta, V., Altintas, M. M., Garborcauskas, G., Stojanovic, N., Winkler, C. A., … Reiser, J. (2017). A tripartite complex of suPAR, APOL1 risk variants and αvβ3 integrin on podocytes mediates chronic kidney disease. Nature Medicine, 23(8), 945–953. 10.1038/nm.4362

Hu, Q., Saleem, K., Pandey, J., Charania, A. N., Zhou, Y., & He, C. (2023). Cell Adhesion Molecules in Fibrotic Diseases. Biomedicines, 11(7), 1995. 10.3390/biomedicines11071995

Hughes, D. C., Wallace, M. A., & Baar, K. (2015). Effects of aging, exercise, and disease on force transfer in skeletal muscle. American Journal of Physiology. Endocrinology and Metabolism, 309(1), E1–E10. 10.1152/ajpendo.00095.2015

Ing-Esteves, S., Kostadinov, D., Marocha, J., Sing, A. D., Joseph, K. S., Laboulaye, M. A., Sanes, J. R., & Lefebvre, J. L. (2018). Combinatorial Effects of Alpha- and Gamma-Protocadherins on Neuronal Survival and Dendritic Self-Avoidance. The Journal of Neuroscience: The Official Journal of the Society for Neuroscience, 38(11), 2713–2729. 10.1523/JNEUROSCI.3035-17.2018

Jabalameli, M. R., Lin, J.-R., Zhang, Q., Wang, Z., Mitra, J., Nguyen, N., Gao, T., Khusidman, M., Sathyan, S., Atzmon, G., Milman, S., Vijg, J., Barzilai, N., & Zhang, Z. D. (2024). Polygenic prediction of human longevity on the supposition of pervasive pleiotropy. Scientific Reports, 14(1), 19981. 10.1038/s41598-024-69069-0

Jacob, J. T., Coulombe, P. A., Kwan, R., & Omary, M. B. (2018). Types I and II Keratin Intermediate Filaments. Cold Spring Harbor Perspectives in Biology, 10(4), a018275. 10.1101/cshperspect.a018275

Kim, Y. J., Cho, M. J., Yu, W. D., Kim, M. J., Kim, S. Y., & Lee, J. H. (2022). Links of Cytoskeletal Integrity with Disease and Aging. Cells, 11(18), 2896. 10.3390/cells11182896

Kircher, M., Witten, D. M., Jain, P., O’Roak, B. J., Cooper, G. M., & Shendure, J. (2014). A general framework for estimating the relative pathogenicity of human genetic variants. Nature Genetics, 46(3), 310–315. 10.1038/ng.2892

Klinkhammer, B. M., Floege, J., & Boor, P. (2018). PDGF in organ fibrosis. Molecular Aspects of Medicine, 62, 44–62. 10.1016/j.mam.2017.11.008

Knaus, B. J., & Grünwald, N. J. (2017). vcfr: A package to manipulate and visualize variant call format data in R. Molecular Ecology Resources, 17(1), 44–53. 10.1111/1755-0998.12549

Kotlyarov, S., & Kotlyarova, A. (2022). Clinical Significance of Lipid Transport Function of ABC Transporters in the Innate Immune System. Membranes, 12(11), 1083. 10.3390/membranes12111083

Lampugnani, M. G., Dejana, E., & Giampietro, C. (2017). Corrigendum: Vascular Endothelial (VE)-Cadherin, Endothelial Adherens Junctions, and Vascular Disease. Cold Spring Harbor Perspectives in Biology, 9(10), a033720. 10.1101/cshperspect.a033720

Lee, M., Cronin, K. A., Gail, M. H., & Feuer, E. J. (2012). Predicting the absolute risk of dying from colorectal cancer and from other causes using population-based cancer registry data. Statistics in Medicine, 31(5), 489–500. 10.1002/sim.4454

Lee, S., Emond, M. J., Bamshad, M. J., Barnes, K. C., Rieder, M. J., Nickerson, D. A., NHLBI GO Exome Sequencing Project—ESP Lung Project Team, Christiani, D. C., Wurfel, M. M., & Lin, X. (2012). Optimal unified approach for rare-variant association testing with application to small-sample case-control whole-exome sequencing studies. American Journal of Human Genetics, 91(2), 224–237. 10.1016/j.ajhg.2012.06.007

Li, J., Patel, V. V., & Radice, G. L. (2006). Dysregulation of cell adhesion proteins and cardiac arrhythmogenesis. Clinical Medicine & Research, 4(1), 42–52. 10.3121/cmr.4.1.42

Li, N., Li, H., Wei, L., Chen, H., Wu, Z., Yuwen, S., & Yang, S. (2024). The Downregulation of MMP23B Facilitates the Suppression of Vitality and Induction of Apoptosis in Endometrial Cancer Cells. Reproductive Sciences, 31(11), 3452–3461. 10.1007/s43032-024-01581-0

Liang, X., Qiu, X., Ma, Y., Xu, W., Chen, S., Zhang, P., Liu, M., & Lin, X. (2023). KRT18 regulates trophoblast cell migration and invasion which are essential for embryo implantation. Reproductive Biology and Endocrinology: RB&E, 21(1), 78. 10.1186/s12958-023-01129-y

Liu, Y., Peng, L., Chen, J., Chen, L., Wu, Y., Cheng, M., Chen, M., Ye, X., & Jin, Y. (2023). EIF5A2 specifically regulates the transcription of aging-related genes in human neuroblastoma cells. BMC Geriatrics, 23(1), 83. 10.1186/s12877-023-03793-6

Magirr, D. (2021). Non-proportional hazards in immuno-oncology: Is an old perspective needed? Pharmaceutical Statistics, 20(3), 512–527. 10.1002/pst.2091

Mak, J. K. L., Qin, C., Krüger, M., Kuukka, A., FinnGen, Hägg, S., Lin, J., & Jylhävä, J. (2025). Large-scale genome-wide analyses with proteomics integration reveal novel loci and biological insights into frailty. Nature Aging, 5(8), 1589–1600. 10.1038/s43587-025-00925-y

Martínez-Férriz, A., Gandía, C., Pardo-Sánchez, J. M., Fathinajafabadi, A., Ferrando, A., & Farràs, R. (2023). Eukaryotic Initiation Factor 5A2 localizes to actively translating ribosomes to promote cancer cell protrusions and invasive capacity. Cell Communication and Signaling: CCS, 21(1), 54. 10.1186/s12964-023-01076-6

McLaren, W., Gil, L., Hunt, S. E., Riat, H. S., Ritchie, G. R. S., Thormann, A., Flicek, P., & Cunningham, F. (2016). The Ensembl Variant Effect Predictor. Genome Biology, 17(1), 122. 10.1186/s13059-016-0974-4

Meng, F.-D., Wei, J.-C., Qu, K., Wang, Z.-X., Wu, Q.-F., Tai, M.-H., Liu, H.-C., Zhang, R.-Y., & Liu, C. (2015). FoxM1 overexpression promotes epithelial-mesenchymal transition and metastasis of hepatocellular carcinoma. World Journal of Gastroenterology, 21(1), 196–213. 10.3748/wjg.v21.i1.196

Ogata, H., Goto, S., Sato, K., Fujibuchi, W., Bono, H., & Kanehisa, M. (1999). KEGG: Kyoto Encyclopedia of Genes and Genomes. Nucleic Acids Research, 27(1), 29–34. 10.1093/nar/27.1.29

Olson, J. D., Schaaf, G. W., Bourland, J. D., & Cline, J. M. (2025). The Wake Forest Nonhuman Primate Radiation Late Effects Cohort. Radiation Research, 204(4), 274–282. 10.1667/RADE-25-00063.1

Pak, K., Uno, H., Kim, D. H., Tian, L., Kane, R. C., Takeuchi, M., Fu, H., Claggett, B., & Wei, L.-J. (2017). Interpretability of Cancer Clinical Trial Results Using Restricted Mean Survival Time as an Alternative to the Hazard Ratio. JAMA Oncology, 3(12), 1692–1696. 10.1001/jamaoncol.2017.2797

Pancho, A., Aerts, T., Mitsogiannis, M. D., & Seuntjens, E. (2020). Protocadherins at the Crossroad of Signaling Pathways. Frontiers in Molecular Neuroscience, 13, 117. 10.3389/fnmol.2020.00117

Pang, X., He, X., Qiu, Z., Zhang, H., Xie, R., Liu, Z., Gu, Y., Zhao, N., Xiang, Q., & Cui, Y. (2023). Targeting integrin pathways: Mechanisms and advances in therapy. Signal Transduction and Targeted Therapy, 8(1), 1. 10.1038/s41392-022-01259-6

Pang, Z., Wang, Y., Ding, N., Chen, X., Yang, Y., Wang, G., Liu, Q., & Du, J. (2019). High PKD2 predicts poor prognosis in lung adenocarcinoma via promoting Epithelial-mesenchymal Transition. Scientific Reports, 9(1), 1324. 10.1038/s41598-018-37285-0

Paukner, M., & Chappell, R. (2021). Window mean survival time. Statistics in Medicine, 40(25), 5521–5533. 10.1002/sim.9138

Paukner, M., Ladner, D. P., & Zhao, L. (2024). Dynamic risk prediction of survival in liver cirrhosis: A comparison of landmarking approaches. PloS One, 19(7), e0306328. 10.1371/journal.pone.0306328

Pays, E. (2020). The Mechanism of Kidney Disease Due to APOL1 Risk Variants. Journal of the American Society of Nephrology: JASN, 31(11), 2502–2505. 10.1681/ASN.2020070954

Peng, S.-Y., Tu, H.-F., Yang, C.-C., Wu, C.-H., Liu, C.-J., Chang, K.-W., & Lin, S.-C. (2018). miR-134 targets PDCD7 to reduce E-cadherin expression and enhance oral cancer progression. International Journal of Cancer, 143(11), 2892–2904. 10.1002/ijc.31638

Prall, T. M., Karl, J. A., Varghese, J. M., Baker, D. A., Minor, N. R., Raveendran, M., Harris, R. A., Rogers, J., Wiseman, R. W., & O’Connor, D. H. (2024). Complete Genomic Assembly of Mauritian Cynomolgus Macaque Killer Ig-like Receptor and Natural Killer Group 2 Haplotypes. Journal of Immunology, 212(11), 1754–1765. 10.4049/jimmunol.2300856

Purcell, S., Neale, B., Todd-Brown, K., Thomas, L., Ferreira, M. A. R., Bender, D., Maller, J., Sklar, P., de Bakker, P. I. W., Daly, M. J., & Sham, P. C. (2007). PLINK: A tool set for whole-genome association and population-based linkage analyses. American Journal of Human Genetics, 81(3), 559–575. 10.1086/519795

Quillen, E. E., Schaaf, G. W., Justice, J. N., Dugan, G. O., Johnson, B., Reed, C., Olson, J. D., & Cline, J. M. (2025). Widespread Multimorbidity in a Cohort of Aging, Radiation-exposed Rhesus Macaques. Radiation Research, 204(4), 283–293. 10.1667/RADE-24-00014.1

R Core Team. (2025). R: A Language and Environment for Statistical Computing. R Foundation for Statistical Computing. https://www.R-project.org/

Sandstedt, J., Vukusic, K., Rekabdar, E., Dellgren, G., Jeppsson, A., Mattsson Hultén, L., & Rotter Sopasakis, V. (2021). Markedly reduced myocardial expression of γ-protocadherins and long non-coding RNAs in patients with heart disease. International Journal of Cardiology, 344, 149–159. 10.1016/j.ijcard.2021.09.046

Schaaf, G. W., Justice, J. N., Quillen, E. E., & Cline, J. M. (2023). Resilience, aging, and response to radiation exposure (RARRE) in nonhuman primates: A resource review. GeroScience, 45(6), 3371–3379. 10.1007/s11357-023-00812-7

Schaaf, G. W., Olson, J. D., Kahn, B. T., Kock, N. D., Caudell, D. L., Lang, R. A., Sills, W. S., Andrews, R. N., Thakur, P., & Cline, J. M. (2025). Postmortem Findings from the Wake Forest University Radiation Late Effects Cohort of Rhesus Monkeys (Macaca mulatta). Radiation Research, 204(4), 294–310. 10.1667/RADE-24-00119.1

Serneels, L., Bammens, L., Zwijsen, A., Tolia, A., Chávez-Gutiérrez, L., & De Strooper, B. (2024). Functional and topological analysis of PSENEN, the fourth subunit of the γ-secretase complex. The Journal of Biological Chemistry, 300(1), 105533. 10.1016/j.jbc.2023.105533

Sivasankar, S., & Xie, B. (2023). Engineering the Interactions of Classical Cadherin Cell-Cell Adhesion Proteins. Journal of Immunology, 211(3), 343–349. 10.4049/jimmunol.2300098

Slowikowski, K. (2016). ggrepel: Automatically Position Non-Overlapping Text Labels with “ggplot2” (p. 0.9.8) [Data set]. 10.32614/CRAN.package.ggrepel

Subramanian, A., Tamayo, P., Mootha, V. K., Mukherjee, S., Ebert, B. L., Gillette, M. A., Paulovich, A., Pomeroy, S. L., Golub, T. R., Lander, E. S., & Mesirov, J. P. (2005). Gene set enrichment analysis: A knowledge-based approach for interpreting genome-wide expression profiles. Proceedings of the National Academy of Sciences of the United States of America, 102(43), 15545–15550. 10.1073/pnas.0506580102

Sundaram, L., Gao, H., Padigepati, S. R., McRae, J. F., Li, Y., Kosmicki, J. A., Fritzilas, N., Hakenberg, J., Dutta, A., Shon, J., Xu, J., Batzoglou, S., Li, X., & Farh, K. K.-H. (2018). Predicting the clinical impact of human mutation with deep neural networks. Nature Genetics, 50(8), 1161–1170. 10.1038/s41588-018-0167-z

Uno, H., Claggett, B., Tian, L., Inoue, E., Gallo, P., Miyata, T., Schrag, D., Takeuchi, M., Uyama, Y., Zhao, L., Skali, H., Solomon, S., Jacobus, S., Hughes, M., Packer, M., & Wei, L.-J. (2014). Moving beyond the hazard ratio in quantifying the between-group difference in survival analysis. Journal of Clinical Oncology: Official Journal of the American Society of Clinical Oncology, 32(22), 2380–2385. 10.1200/JCO.2014.55.2208

Wada, Y., Inoko, M., Ishihara, K., Fukumoto, K., Tsurudome, Y., Horiguchi, M., Fujimura, A., & Ushijima, K. (2025). Aging Reduces ATP-Binding Cassette Transporter Expression in Brain Microvessels of Mice. Pharmaceuticals, 18(2), 191. 10.3390/ph18020191

Wang, Z., Shi, J., Zhang, Y., Luo, Y., Rao, Y., Qu, J., Gai, X., & Sun, Y. (2026). LAMC2 Drives Airway Remodeling in COPD via EMT Regulation Through the AKT Pathway. International Journal of Chronic Obstructive Pulmonary Disease, 21, 580964. 10.2147/COPD.S580964

Wickham, H. (2016). ggplot2: Elegant Graphics for Data Analysis (2nd ed. 2016). Springer International Publishing : Imprint: Springer. 10.1007/978-3-319-24277-4

Wickham, H., Averick, M., Bryan, J., Chang, W., McGowan, L., François, R., Grolemund, G., Hayes, A., Henry, L., Hester, J., Kuhn, M., Pedersen, T., Miller, E., Bache, S., Müller, K., Ooms, J., Robinson, D., Seidel, D., Spinu, V., … Yutani, H. (2019). Welcome to the Tidyverse. Journal of Open Source Software, 4(43), 1686. 10.21105/joss.01686

Yang, Z., Wu, H., Hou, Y., Yuan, H., & Chen, Z. (2021). Dynamic prediction and analysis based on restricted mean survival time in survival analysis with nonproportional hazards. Computer Methods and Programs in Biomedicine, 207, 106155. 10.1016/j.cmpb.2021.106155

Yu, G., Wang, L.-G., Han, Y., & He, Q.-Y. (2012). clusterProfiler: An R package for comparing biological themes among gene clusters. Omics: A Journal of Integrative Biology, 16(5), 284–287. 10.1089/omi.2011.0118

Zhang, X., Miao, Y., Yu, X., Zhang, Y., Jiang, G., Liu, Y., Yu, J., Han, Q., Zhao, H., & Wang, E. (2015). C6orf106 enhances NSCLC cell invasion by upregulating vimentin, and downregulating E-cadherin and P120ctn. Tumour Biology: The Journal of the International Society for Oncodevelopmental Biology and Medicine, 36(8), 5979–5985. 10.1007/s13277-015-3274-9

Zhao, L., Claggett, B., Tian, L., Uno, H., Pfeffer, M. A., Solomon, S. D., Trippa, L., & Wei, L. J. (2016). On the restricted mean survival time curve in survival analysis. Biometrics, 72(1), 215–221. 10.1111/biom.12384

Zhao, L., Murray, S., Mariani, L. H., & Ju, W. (2020). Incorporating longitudinal biomarkers for dynamic risk prediction in the era of big data: A pseudo-observation approach. Statistics in Medicine, 39(26), 3685–3699. 10.1002/sim.8687

Zheng, Y., Li, P., Huang, H., Ye, X., Chen, W., Xu, G., & Zhang, F. (2021). Androgen receptor regulates eIF5A2 expression and promotes prostate cancer metastasis via EMT. Cell Death Discovery, 7(1), 373. 10.1038/s41420-021-00764-x

Zhou, X., & Stephens, M. (2012). Genome-wide efficient mixed-model analysis for association studies. Nature Genetics, 44(7), 821–824. 10.1038/ng.2310

Zhuo, Y., Zhang, J., Laboy, M., & Lasky, J. A. (2004). Modulation of PDGF-C and PDGF-D expression during bleomycin-induced lung fibrosis. American Journal of Physiology. Lung Cellular and Molecular Physiology, 286(1), L182–188. 10.1152/ajplung.00083.2003

