## Supplementary Table 1 for "Pseudo-observations of longevity improve detection of resilience-associated genes in the Radiation Late Effects Cohort of rhesus macaques"

**Supplementary Table 1: Summary of significant longevity-associated variants**

| Gene | Ensembl Gene ID | Rhesus SNV | p-value | beta | se | af | Consequence Detail | Ensemble Feature | CADD Raw | CADD PHRED |
|---|---|---|---|---|---|---|---|---|---|---|
| **KCNK15** | ENSG00000124249 | 10:79992432:C:G | 2.73E-14 | -1.042 | 0.128 | 0.220 | synonymous | CTCF Binding Site | 0.368 | 5.069 |
| | ENSG00000124249 | 10:79992345:T:G | 5.74E-13 | -0.919 | 0.121 | 0.323 | synonymous | CTCF Binding Site | 0.118 | 2.254 |
| **KRT18** | ENSG00000111057 | 11:52512929:C:A | 3.99E-14 | 2.712 | 0.338 | 0.449 | synonymous | Promoter | 1.519 | 15.86 |
| **ANKRD36C** | ENSG00000174501 | 13:16735386:A:G | 5.49E-14 | -0.937 | 0.118 | 0.297 | missense | NA | -0.810 | 0.024 |
| **ESYT2** | ENSG00000117868 | 3:184778107:A:G | 1.01E-13 | -1.038 | 0.132 | 0.199 | missense | Promoter | 0.286 | 4.125 |
| | ENSG00000117868 | 3:184778104:A:G | 5.91E-10 | -0.796 | 0.124 | 0.353 | missense | Promoter | 0.105 | 2.134 |
| **PRR20B** | ENSG00000204918 | 17:35777979:A:C | 5.10E-13 | -0.924 | 0.121 | 0.314 | synonymous | NA | 0.525 | 6.764 |
| **CAMKMT** | ENSG00000143919 | 13:64230251:A:G | 5.61E-13 | 0.943 | 0.124 | 0.637 | intron | NA | 0.102 | 2.108 |
| **SPTBN4** | ENSG00000160460 | 19:39737253:A:G | 1.02E-12 | -1.027 | 0.137 | 0.190 | synonymous | NA | 0.012 | 1.41 |
| **PSG11** | ENSG00000243130 | 19:43060890:A:G | 1.77E-12 | 2.315 | 0.309 | 0.434 | synonymous | NA | -0.079 | 0.918 |
| | ENSG00000243130 | 19:43064491:G:A | 8.70E-10 | 2.372 | 0.373 | 0.462 | synonymous | NA | 0.048 | 1.663 |
| **NPIPA3** | ENSG00000224712 | 20:14708913:A:G | 2.87E-12 | 0.915 | 0.124 | 0.632 | missense | Promoter Flanking Region | 0.270 | 3.936 |
| **FOXM1** | ENSG00000111206 | 11:2963786:G:A | 4.74E-12 | -1.218 | 0.168 | 0.107 | synonymous | NA | 0.957 | 11.08 |
| **PSG4** | ENSG00000243137 | 19:43323582:C:G | 6.27E-12 | -0.876 | 0.121 | 0.316 | missense | NA | 0.026 | 1.507 |
| **IGFBPL1** | ENSG00000137142 | 15:45686948:G:A | 7.69E-12 | -0.903 | 0.126 | 0.239 | synonymous | Promoter | 0.248 | 3.677 |
| **EIF5A2** | ENSG00000163577 | 2:75807718:A:G,C | 8.64E-12 | 1.601 | 0.224 | 0.363 | missense | Promoter | 2.961 | 23.3 |
| | ENSG00000163577 | 2:75807711:G:T | 2.80E-10 | 1.514 | 0.230 | 0.368 | missense | Promoter | 3.276 | 24.1 |
| | ENSG00000163577 | 2:75807705:A:G,C | 1.20E-09 | 1.454 | 0.230 | 0.367 | synonymous | Promoter | 1.084 | 12.61 |
| **ADRA2A** | ENSG00000150594 | 9:111638375:T:G | 1.35E-11 | -1.104 | 0.156 | 0.139 | missense | Promoter | 2.206 | 20.9 |
| **BICRA** | ENSG00000063169 | 19:47714463:T:C | 3.00E-11 | -1.199 | 0.173 | 0.107 | missense | NA | 2.797 | 23 |
| **ZNF426** | ENSG00000130818 | 19:9322159:T:C | 3.21E-11 | -1.188 | 0.171 | 0.115 | missense | NA | 1.002 | 11.65 |
| **C9orf85** | ENSG00000155621 | 15:87347683:A:G | 3.97E-11 | -0.852 | 0.124 | 0.280 | intron | NA | -0.032 | 1.149 |
| **C4orf48** | ENSG00000243449 | 5:2111247:T:C | 5.71E-11 | -0.904 | 0.132 | 0.222 | synonymous | Promoter | 1.316 | 14.5 |
| **RASIP1** | ENSG00000105538 | 19:48721643:A:G | 6.54E-11 | -1.586 | 0.233 | 0.051 | synonymous | CTCF Binding Site | 0.521 | 6.72 |
| **KBTBD11** | ENSG00000176595 | 8:2513372:C:G | 7.30E-11 | -0.965 | 0.141 | 0.256 | synonymous | NA | 0.940 | 10.88 |
| **ILRUN** | ENSG00000196821 | 4:134843174:A:G | 1.18E-10 | 2.223 | 0.330 | 0.442 | missense | NA | 2.372 | 22 |
| | ENSG00000196821 | 4:134843229:T:C | 1.90E-10 | 1.798 | 0.269 | 0.410 | missense | NA | 4.052 | 27.4 |
| **AC073957.1** | ENSG00000225146 | 3:34098771:G:A | 1.37E-10 | -0.852 | 0.127 | 0.274 | upstream | Promoter | 1.585 | 16.28 |

| | | | | | | | | | | |
|---|---|---|---|---|---|---|---|---|---|---|
| **ZC3HAV1** | ENSG00000105939 | 3:165467290:C:T | 1.81E-10 | -1.096 | 0.164 | 0.135 | synonymous | NA | 0.047 | 1.656 |
| **APOL1** | ENSG00000100342 | 10:15309639:T:C | 3.45E-10 | -0.823 | 0.126 | 0.316 | missense | NA | -0.808 | 0.024 |
| **PTGER1** | ENSG00000160951 | 19:14093916:G:C | 3.45E-10 | -0.944 | 0.144 | 0.212 | synonymous | Promoter | 0.266 | 3.894 |
| **PDCD7** | ENSG00000090470 | 7:41661986:A:G | 5.21E-10 | -0.911 | 0.140 | 0.222 | missense | Promoter | 1.013 | 11.78 |
| **MIA3** | ENSG00000154305 | 1:80306926:G:A | 5.28E-10 | -1.835 | 0.284 | 0.062 | splice,synonymous | NA | 0.593 | 7.444 |
| **PKD2** | ENSG00000118762 | 5:86399759:C:T | 1.01E-09 | -0.878 | 0.138 | 0.171 | missense | Promoter | 2.062 | 19.73 |
| **ASTN2** | ENSG00000148219 | 15:25402984:C:G | 1.18E-09 | -0.928 | 0.147 | 0.205 | synonymous | Promoter | 0.771 | 9.107 |
| **TIGD1** | ENSG00000221944 | 12:119972921:C:T | 2.35E-09 | -0.868 | 0.140 | 0.295 | synonymous | NA | 0.560 | 7.121 |
| **COL23A1** | ENSG00000050767 | 6:175928735:T:C | 3.09E-09 | -0.772 | 0.125 | 0.393 | missense | Promoter | 1.899 | 18.48 |
| **MMP23B** | ENSG00000189409 | 1:222764801:G:C | 3.89E-09 | -1.141 | 0.187 | 0.090 | missense | Promoter | 2.369 | 21.9 |
| **EMILIN2** | ENSG00000132205 | 18:64143664:T:C | 7.49E-09 | -0.914 | 0.153 | 0.156 | synonymous | Promoter | -0.069 | 0.962 |
| **HES4** | ENSG00000188290 | 1:223421865:G:C | 8.75E-09 | -0.909 | 0.153 | 0.162 | missense | Promoter | 1.317 | 14.51 |
| **GOLGA8K** | ENSG00000249931 | 7:7372273:C:G | 8.83E-09 | -0.790 | 0.133 | 0.280 | synonymous | NA | 0.161 | 2.691 |
