## Supplementary Table 2 for "Pseudo-observations of longevity improve detection of resilience-associated genes in the Radiation Late Effects Cohort of rhesus macaques"

**Supplementary Table 2: Total Burden Result Statistics from Cadherin Combined GSEA Method Leading-Edge Genes**

| Pathway | Fibrosis Phenotype | Animal Group | Weighing Scheme | p-value | Beta | SE | $R^2$ |
|---|---|---|---|---|---|---|---|
| **Cadherin** | Pulmonary | all | threshold | **0.0500** | 0.0163 | 0.0082 | 0.0421 |
| | | | linear | 0.0511 | 0.0217 | 0.0110 | 0.0418 |
| | | | unweighted | 0.0676 | 0.0015 | 0.0008 | 0.0381 |
| | | irradiated | threshold | **0.0347** | 0.0202 | 0.0094 | 0.0452 |
| | | | linear | **0.0371** | 0.0274 | 0.0130 | 0.0440 |
| | | | unweighted | 0.1079 | 0.0016 | 0.0010 | 0.0264 |
| | Myocardial | all | threshold | 0.0720 | 0.0140 | 0.0077 | 0.1720 |
| | | | linear | **0.0378** | 0.0216 | 0.0103 | 0.1794 |
| | | | unweighted | 0.1323 | 0.0012 | 0.0008 | 0.1653 |
| | | irradiated | threshold | 0.1535 | 0.0124 | 0.0086 | 0.1228 |
| | | | linear | 0.1294 | 0.0180 | 0.0118 | 0.1252 |
| | | | unweighted | 0.3785 | 0.0008 | 0.0009 | 0.1113 |
